# Performance of large scale pooled CRISPR screens is dependent on Cas9 expression levels

**DOI:** 10.1101/2021.07.13.452178

**Authors:** João M. Fernandes Neto, Cor Lieftink, Katarzyna Jastrzebski, Lenno Krenning, Matheus Dias, Ben Morris, Daimy van der Ven, Hester Heimans, René H. Medema, René Bernards, Roderick L. Beijersbergen

**Author notes:** Corresponding authors: João Neto, Division of Molecular Carcinogenesis, The Netherlands Cancer Institute, Plesmanlaan 121, 1066 CX Amsterdam, The Netherlands; Roderick Beijersbergen, Division of Molecular Carcinogenesis, The Netherlands Cancer Institute, Plesmanlaan 121, 1066 CX Amsterdam, The Netherlands. These authors contributed equally to this work (Cor Lieftink, Katarzyna Jastrzebski).

## Abstract

**Background:** The widespread application of CRISPR/Cas9 technology has yielded numerous findings in biomedical research in recent years, making it an invaluable tool for gene knockout and for high-throughput screening studies. In (low-throughput) gene knockout studies, editing efficiency is not a major concern because only a few edited clones are necessary for a successful assay. However, in large scale pooled screening studies, editing efficiency is a major concern because each sgRNA has to knockout its target gene in a large cell population in a short period of time. Therefore, a thorough understanding of the role that key factors play in determining CRISPR knockout efficiency is essential to improve the performance of pooled CRISPR screening.

**Methods:** In this study, cell lines with different expression levels of CAS9 were generated and used to determine gene-editing efficiency. Collections of sgRNAs targeting essential genes were used to study their depletion in the different cell line models.

**Results:** Using cell lines with variable expression of Cas9, we confirmed that editing efficiency and speed are mostly dependent on the sgRNA sequence and Cas9 expression, respectively. Importantly, we show that the strategy employed for delivering sgRNAs and Cas9 to cells impacts the performance of high-throughput screens, which is improved in conditions with higher Cas9 expression.

**Conclusions:** Our findings highlight the importance of optimizing Cas9 expression levels when performing gene editing experiments and provide guidance on the necessary decisions for implementing optimal pooled CRISPR screening strategies.

## INTRODUCTION

The simplicity, speed and low-cost of CRISPR technology has led to its widespread application in biomedical research in recent years. In particular, large scale pooled CRISPR screening technologies have yielded significant discoveries in a variety of research fields (1–4). Since its establishment, the technology has undergone many improvements. The largest one, arguably, is short guide RNA (sgRNA) sequence design which has improved to a level where efficient genome-wide screening is possible with sgRNA libraries containing only two independent sgRNAs per gene (5). The CRISPR toolbox has been expanded with many different vector systems and versions of Cas9, most of them easily available through non-profit plasmid repository Addgene. These systems differ in the types of promoter (constitutive or inducible), the presence of fusion transcripts (with selectable markers or fluorescent proteins) and the presence or absence of the sgRNA expression cassette. Although beneficial, the availability of many different constructs can also complicate the selection of an optimal CRISPR screening platform and strategy. Thus, understanding which factors influence CRISPR editing efficiency is essential to optimize the performance of high-throughput pooled CRISPR screening.

In mammalian cells, gene editing using the CRISPR/Cas9 system is a multistep process that can take several days, being mostly dependent on sgRNA sequence and Cas9 expression (6). When the experimental goal is to knockout a single gene, several clones are usually screened until a few that are successfully edited are identified. In this low-throughput setting time is not a major issue. However, when the goal is to knockout many genes (or even the whole genome) in a large population of cells – as is the case for large scale pooled functional genetic screens – time is of the essence. By increasing the number of cells that are effectively edited when a selective pressure is applied, we can reduce screening noise. But if the editing is slow, it may take weeks to get to a point where the majority of the cells are effectively edited, which increases the logistical hurdles of culturing such a large number of cells.

In recent years, several studies have found that by increasing Cas9 expression (through the use of Cas9 RNPs, optimized promoters or simply by increasing the MOI of the vector which expresses Cas9) the editing speed also increased (6–9). Therefore, we hypothesized that in a pooled CRISPR screen, by optimizing Cas9 expression, we might reduce the time necessary to achieve editing in the majority of the cell population, thereby reducing logistical hurdles and potentially improving screening quality.

We reasoned that sgRNA sequence will have a major impact on the editing efficiency of both low- and high-throughput settings. However, in high-throughput screens multiple sgRNAs targeting the same gene are in general present, therefore the effect of a less active or even inactive sgRNA can be mitigated on the gene level. Additionally, sgRNA design has improved a lot in the past years and most commercially available sgRNA libraries perform well.

A critical aspect of pooled CRISPR screens is the introduction of a single (unique) sgRNA in each cell; another is the sgRNA library representation, meaning that each sgRNA will be present in (typically) 200-250 different cells. To achieve a single integration of each sgRNA, the target cells are transduced with the lentiviral vector used to deliver the sgRNA using a low multiplicity of infection (MOI). Because the site of integration of the lentivirus will likely be different in each cell, the transduced cells will have variable expression levels of the integrated construct. Because of this variation, and building on the knowledge that higher Cas9 expression yields faster knockouts, it is reasonable to hypothesize that it is not ideal to transduce cells with a lentiviral backbone expressing both the Cas9 and the sgRNA on the same vector. This could produce a large range of Cas9 levels across the population of cells leading to a situation where, for example, two different cells carrying the same sgRNA would express different levels of Cas9. This would lead to varying editing efficiencies and thereby to inferior screen performance. On the other hand, by first transducing a construct expressing only Cas9 into cells, followed by selection and expansion of a population with homogeneous high levels of Cas9 expression and transduction with the sgRNA library, the whole population of cells should achieve gene knockouts faster. Additionally, by splitting the Cas9 and the sgRNA expression into 2 different lentiviral vectors, higher viral titers can be achieved, especially in the construct which expresses the sgRNA library – where high titers are a necessity (10). Most publicly available sgRNA libraries can be purchased in a “1-vector system” which delivers both the sgRNA library and Cas9 expression at the same time, or in a “2-vector system” which requires first a vector to deliver Cas9 to the cells followed by a second vector to deliver the sgRNA library. Despite its relevance to the CRISPR screening field, to our knowledge, no comprehensive comparison has been made to address the differences between these two systems in screening performance until now.

In this study, we investigated the influence of sgRNA sequences and Cas9 expression on CRISPR/Cas9-mediated gene editing. Our data demonstrate that screen performance improves with higher editing speed, achieved by increased Cas9 expression levels. We show that high Cas9 expression levels can easily be achieved by choosing the right strategy for delivering Cas9 to cells.

## RESULTS

### Editing efficiency is dependent on sgRNA sequence and Cas9 expression

To assess the effect of different Cas9 expression levels on editing efficiency over time, we transduced SW480 cells with Lenti-iCas9-neo, an inducible lentiviral vector where Cas9 is fused to a 2A self-cleaving peptide followed by EGFP. With this system we could induce different levels of Cas9 expression by titration of doxycycline. Cas9 expression was monitored by analyzing GFP levels by flow cytometry (Fig. 1A and Suppl. Fig. 1A). This cell line was named “SW480_iCas9”. We determined that culturing SW480_iCas9 cells with 10, 40 and 1000 ng/mL of doxycycline induced low, medium and high levels of Cas9 expression, respectively. Cas9 expression levels were confirmed by Western blot (Fig. 1B) and by flow cytometry (Fig. 1C and Suppl. Fig. 1B).

**Figure 1:**
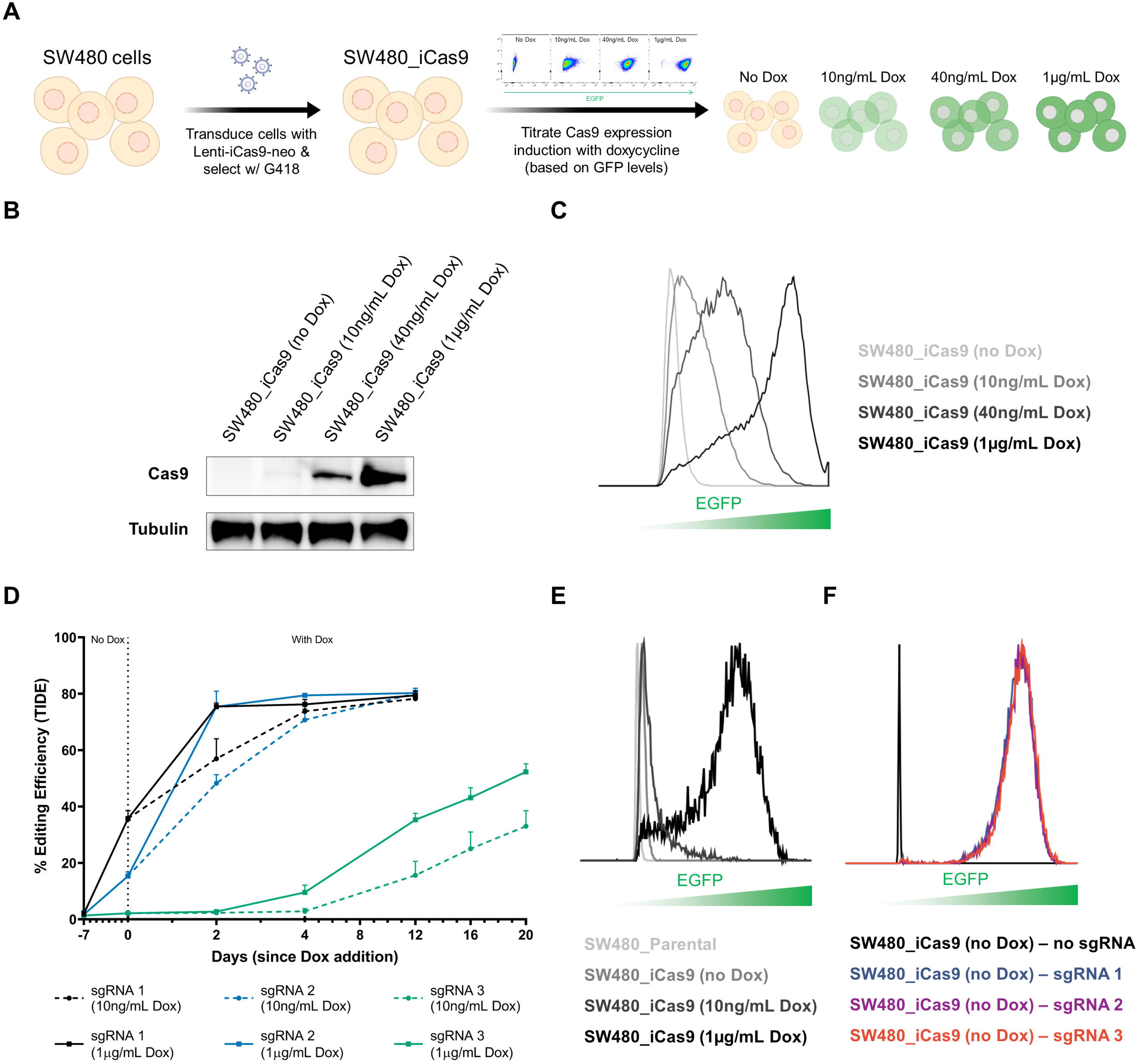
Editing efficiency and speed are dependent on the sgRNA sequence and Cas9 expression. **A,** Schematic of the generation of the SW480_iCas9 cells. Flow cytometry was used to determine the doxycycline concentration necessary to induce different levels of Cas9 expression (low, medium or high), based on EGFP expression. **B,** Analysis of the levels of Cas9 expression using different concentrations of doxycycline by western blot. Tubulin was used as loading control. **C,** Analysis of the levels of Cas9 expression using different concentrations of doxycycline. The level of Cas9 expression was assessed (indirectly) by examining EGFP levels (x-axis) by flow cytometry. **D,** Analysis of editing efficiency in relation to sgRNA sequence and Cas9 expression levels. SW480_iCas9 cells were transduced with 3 different sgRNAs and selected on puromycin for 7 days. At day 7, a reference sample was taken, and remaining cells were treated with either 10 ng/mL or 1 μg/mL doxycycline. Cells were harvested at the indicated time-points. Gene editing efficiency was determined using TIDE analysis. **E,** Analysis of the levels of Cas9 expression in parental SW480 cells and in SW480_iCas9 cells (using different concentrations of doxycycline). Cells were harvested at day 13 of the experiment described in Fig. 1D. The level of Cas9 expression was assessed (indirectly) by examining EGFP levels (x-axis) by flow cytometry. **F,** Analysis of sgRNA-GFP expression in SW480_iCas9 cells. Before the start of the experiment described in Fig. 1D, the level of sgRNA expression was assessed (indirectly) by examining EGFP levels (x-axis) by flow cytometry, to confirm equal infection efficiency and thereby similar sgRNA expression levels.

To assess the effect of sgRNA sequence on editing efficiency over time, we cloned 3 sgRNAs (each targeting a different location in the genome) into lentiGuide-Puro- T2A-GFP (Suppl. Fig. 1C) and transduced them into SW480_iCas9 cells. As illustrated in Suppl. Fig. 1D, we measured the editing efficiency achieved by the 3 different sgRNAs, before and after exposure to 10ng/mL or 1µg/mL doxycycline at different time-points, using TIDE analysis (11) (Fig. 1D). We observed a large difference in editing efficiency across the three sgRNAs, especially between sgRNA 3 and the other two. Further, we observed that editing occurred faster in the cells cultured with 1µg/mL doxycycline. This data confirms previous reports indicating that editing efficiency and editing speed are highly dependent on the sgRNA sequence and Cas9 expression, respectively (6,8,12,13). We noticed that in the cells transduced with the most efficient sgRNAs, a large percentage of cells were already edited before the addition of doxycycline to the cells due to a minimal level of Cas9 expression even in the absence of doxycycline (Fig. 1E and Suppl. Fig.1E). Importantly, this low expression was enough to drive the editing of almost 40% of the cell population transduced with the best sgRNA, in just 7 days. This indicates that in the presence of a very good sgRNA minimal Cas9 expression is sufficient to induce editing. However, in a large scale sgRNA library setting, it is unlikely that all sgRNAs will behave like sgRNA 1. To ensure faster editing even with less effective sgRNAs, cells with a high level of Cas9 expression should be used. Of note, all 3 sgRNAs had equal transduction efficiencies (Fig. 1F), indicating equal number of integrations and similar sgRNA expression levels in the cells, which excludes the possibility that the observed differences in editing efficiencies between the 3 sgRNAs could be due to different levels of sgRNA expression.

### Performance of pooled CRISPR screens is improved in conditions with higher Cas9 expression

The performance of high-throughput loss-of-function (drop-out) screens is commonly evaluated by the ability of a sgRNA library to distinguish essential from non-essential genes (14,15); in other words, a strong depletion of sgRNAs targeting essential genes compared to non-essential genes is an indication of good screen performance. To assess the effect of Cas9 expression on the performance of large scale CRISPR screens, we generated a custom sgRNA library targeting essential genes and safe-havens (called “E/SH library”) and performed drop-out screens (Fig. 2A) in SW480_iCas9 cells cultured with different concentrations of doxycycline and hence expressing different levels of Cas9 (Fig. 2A). The E/SH library was comprised of 696 sgRNAs – 486 sgRNAs targeting a set of 46 essential genes (with on average 10 sgRNAs per gene) and 210 sgRNAs targeting safe-haven regions, as negative controls (Supplementary Table 1). Analysis of the screens showed that with higher Cas9 expression, a greater separation between the positive and negative controls was observed (Fig. 2B and Supplementary Table 1). This indicates that cells with higher Cas9 expression display more rapid gene editing, allowing for longer selection and thus larger depletion, which is beneficial to the screen’s performance.

**Figure 2:**
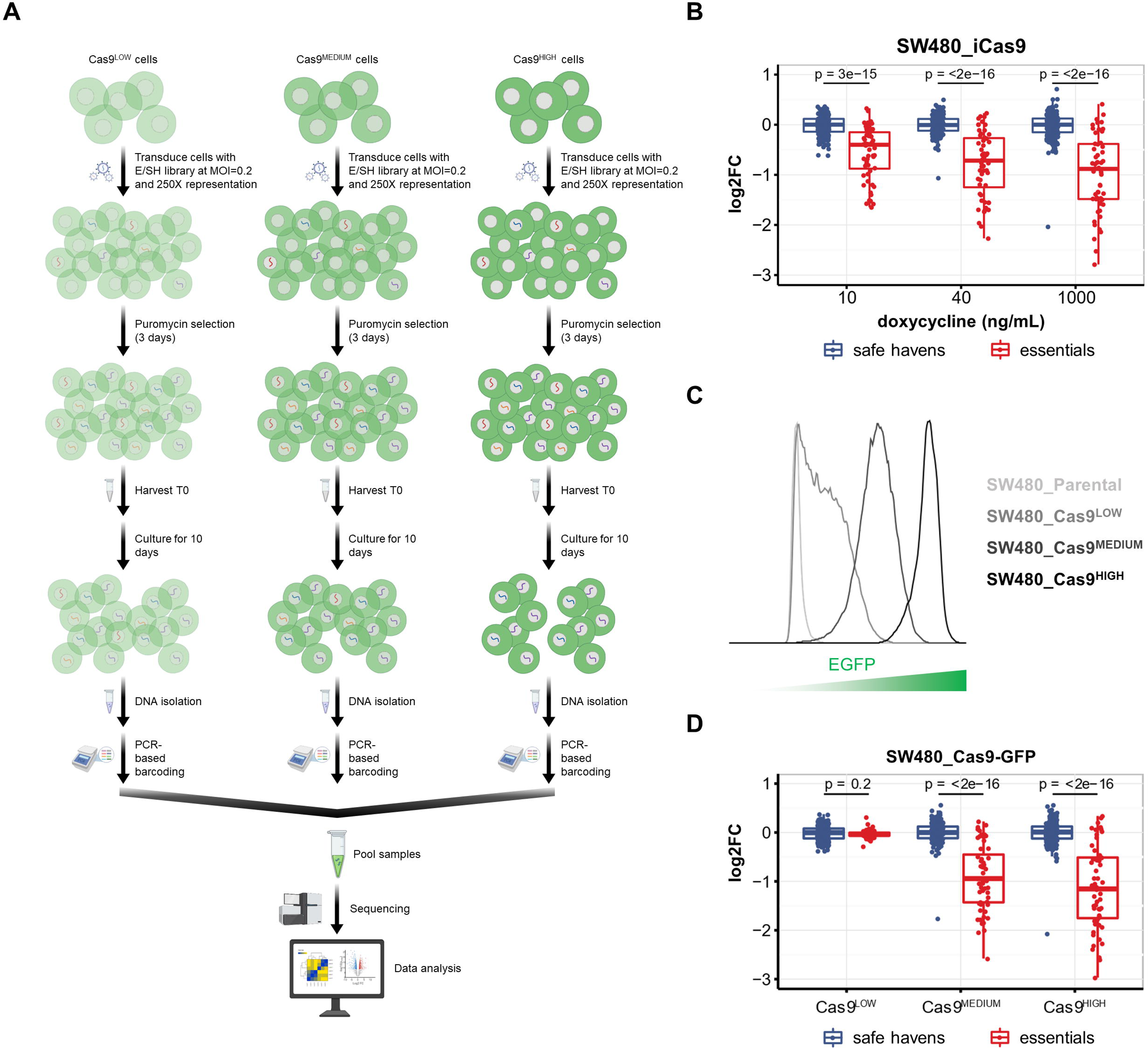
Performance of pooled CRISPR screens is improved in conditions with higher Cas9 expression. **A,** Schematic of the different high-throughput screens performed in cells expressing low, medium and high levels of Cas9. **B,** Analysis of the of the depletion (log2 Fold Change) of the sgRNAs targeting essential genes and safe-havens in SW480_iCas9 cells with different Cas9 expression levels. SW480_iCas9 cells were screened with a library of sgRNAs targeting essential genes and safe- havens. Cells were treated with 10, 40 or 1000 ng/mL of doxycycline and cultured for approximately 8 population doublings. Box plot shows the median, interquartile range and the smallest and largest values no more than 1.5 times the interquartile range. Comparisons were made using the Wilcoxon test. **C,** Analysis of the levels of Cas9 expression in SW480_Cas9-GFP cells. The level of Cas9 expression was assessed (indirectly) by examining GFP levels (x-axis) by flow cytometry. **D,** Analysis of the of the depletion (log2 Fold Change) of the sgRNAs targeting essential genes and safe-havens in SW480_Cas9-GFP cells with different Cas9 expression levels. Cells were screened with a library of sgRNAs targeting essential genes and safe-havens and cultured for approximately 8 population doublings. Box plot shows the median, interquartile range and the smallest and largest values no more than 1.5 times the interquartile range. Comparisons were made using the Wilcoxon test.

To confirm that the observed correlation between screen performance and level of Cas9 expression was independent of cell line and Cas9 expression vector, we expressed Cas9 in 3 cell lines (SW480, A375 and HEK 293T) using lentiCas9-EGFP – a constitutive lentiviral vector where Cas9 is fused to a 2A self-cleaving peptide followed by EGFP. With this system, we could easily generate cells with low, medium and high levels of Cas9 expression by sorting cells with low, medium and high GFP expression, respectively (Fig. 2C and Suppl. Fig. 2A-C). We then performed drop-out screens in these cell lines with the E/SH library. Analysis of the screens showed that, independent of cell line, the depletion of the sgRNAs targeting essential genes was proportional to the level of Cas9 expression in the cells, confirming our hypothesis (Fig. 2D, Suppl. Fig. 2D-E and Supplementary Table 1).

### 2-vector CRISPR system improves the performance of pooled CRISPR screens

CRISPR/Cas9 is a two-component system which requires a sgRNA sequence to direct the Cas9 endonuclease to the target site. In large scale pooled CRISPR screens, to prevent the presence of more than one sgRNA per cell, cells have to be transduced with a low MOI (usually around 0,3). When using a “1-vector system”, most infected cells will integrate one copy of a vector encoding the sgRNA and Cas9. Depending on the position of the integration, which is unpredictable, this could result in varying levels of sgRNA and Cas9 expression across a population of cells. When using a “2-vector system”, a first vector is used to express Cas9, usually at high MOI and then a second vector is used to introduce the sgRNA library, at low MOI. This way, although the expression of sgRNAs will still vary depending on the integration site, the level of Cas9 expression can be controlled/optimized.

To assess whether the strategy employed for delivering sgRNAs and Cas9 to cells impacts the performance of high-throughput screens, we screened MCF10A cells using both a 1-vector and a 2-vector system. For both screening systems, the same collection of sgRNAs (Brunello library) was used. In the 1-vector system screen, the sgRNA library and Cas9 were introduced using the lentiCRISPR v2 version of the Brunello library, using a low MOI. In the 2-vector system screen, we first transduced MCF10A cells with Edit-R – an inducible lentiviral vector to express Cas9 – and selected a clone with high Cas9 expression upon doxycycline treatment and undetectable Cas9 expression in the absence of doxycycline. Then we transduced cells with the lentiGuide-Puro version of the Brunello library, using a low MOI (Fig. 3A and Suppl. Fig. 3A-C). Importantly, we observed that Cas9 expression was higher in the cells for which the 2-vector approach was used (Fig. 3B). To evaluate the performance of each screen, we used a set of sgRNAs targeting 50 essential and 50 non-essential genes, with 4 sgRNAs per gene. This set of genes has been used to benchmark the performance of different technologies perturbing gene expression (14,16). Not surprisingly, we observed significantly more depletion of sgRNAs targeting essential genes in the screen where the 2-vector approach was used, confirming that increasing the level of Cas9 expression improved screen efficiency and indicating that the strategy employed for delivering sgRNAs and Cas9 to cells impacts the performance of high-throughput screens (Fig. 3C and Supplemental Table 2). Of note, ROC curves show a clear benefit of the 2-vector system for individual sgRNAs but on the gene level, using RRA to generate a gene depletion score, the benefit is limited (Fig. 3D-E).

**Figure 3:**
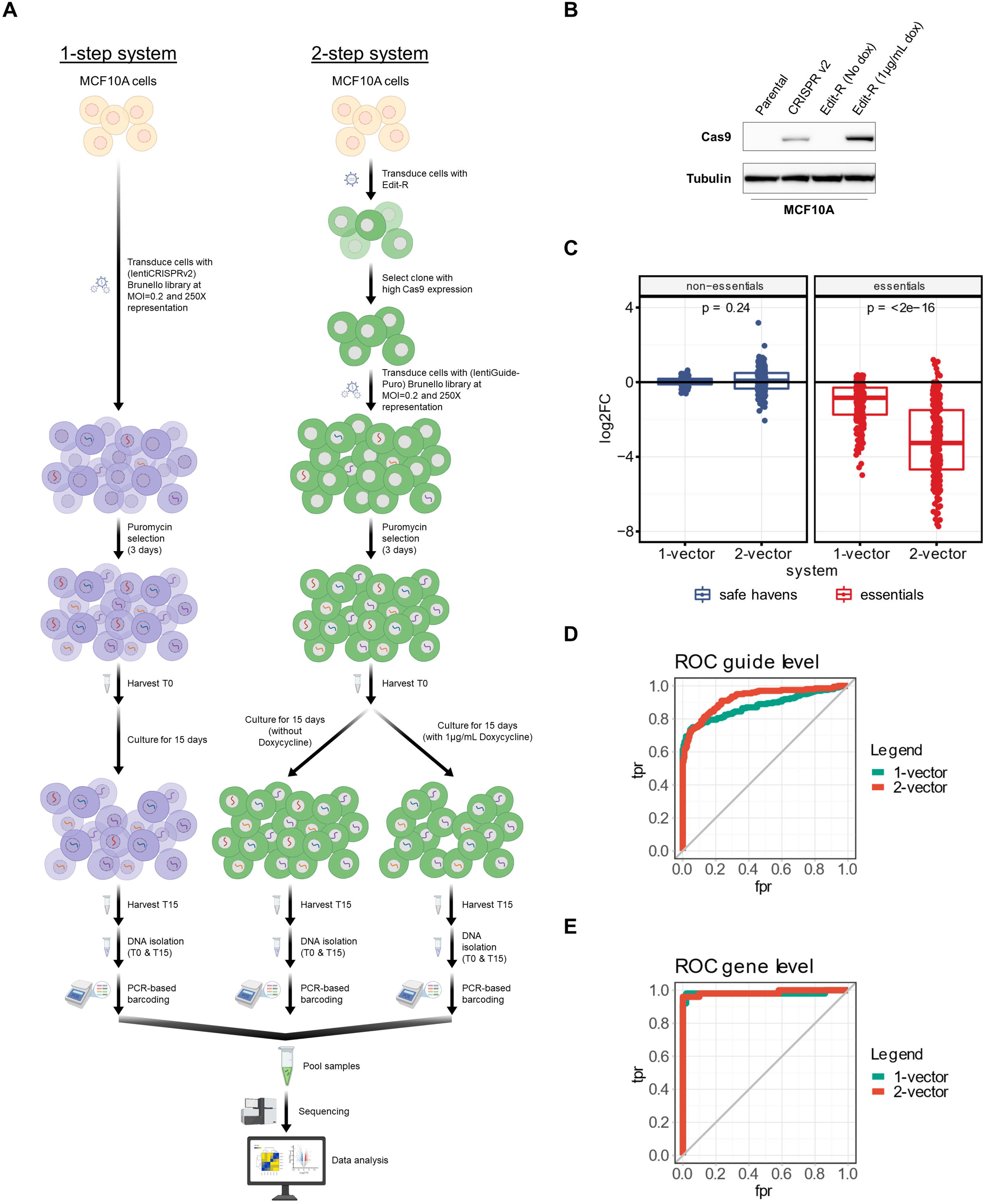
Using a two-vector CRISPR system improves the performance of pooled CRISPR screens. **A,** Schematic of the screen setup using the 1- and 2-step systems. **B,** Comparison of the Cas9 expression induced by a 1-step and by a 2-step system. MCF10A_CRISPR v2 cells were transduced with the lentiCRISPR v2 version of the Brunello library using a MOI of 0,3 and MCF10A_Edit-R cells were transduced with Edit-R using a MOI >1. The level of Cas9 expression was assessed by examining Cas9 levels in the western blot. Tubulin was used as loading control. **C,** Analysis of the depletion (log2 Fold Change) of the sgRNAs targeting essential and non-essential genes. MCF10A cells were screened with the same genome-wide sgRNA library (Brunello), using either a 1-step or a 2-step system. Box plot shows the median (horizontal line), interquartile range (hinges), and the smallest and largest values no more than 1.5 times the interquartile range (whiskers). Comparisons were made using the Wilcoxon test. **D,** ROC curves for individual sgRNAs based on the DESeq2 results sorted on the DESeq2 statistic in increasing order of p-values for essential genes. **E,** ROC curves for genes based on the rank column in the RRA output, in increasing order of p-values for essential genes. FPR, false-positive rate; TPR, true-positive rate.

## DISCUSSION

In this study, we addressed an important and outstanding issue for large scale pooled CRISPR screening – the influence of sgRNA sequence and Cas9 expression on the performance of pooled CRISPR screens. Our findings suggest that in a heterogeneous population of cells, gene editing via CRISPR/Cas9 behaves similarly to an enzymatic reaction, i.e. the percentage of edited cells increases over time until it reaches a plateau. Although the editing rate depends on factors such as sgRNA design (specificity of target site selection), target region (open versus closed chromatin) and DNA repair capacity (NHEJ versus HDR) (17), our data show that editing speed is highly dependent on the level of Cas9 expression, while the maximum achievable editing efficiency is mostly dependent on sgRNA sequence (Fig. 1D). In resistance screens in particular, it is desirable that the screening cells have completed the creation of knock-outs followed by reduction of protein levels before drug treatments are initiated, in order to increase the efficiency to recover resistant cells in the population; if the editing process is slow, it may take weeks to reach that point, which increases the logistical hurdles of culturing such a large number of cells. Besides the logistical advantages, our data show that increasing the editing speed also improves the screen’s performance. Because of that, when performing a pooled CRISPR screen, ensuring high levels of Cas9 expression is essential. Nowadays, there are many different vectors available to express Cas9 and choosing the wrong one can significantly affect the screen’s performance. We show that the use of a 2-vector system, which allows for the generation of a homogenous, high Cas9-expressing cell population, is preferred. While there are many vectors to introduce Cas9 expression available, we recommend the use of a construct where Cas9 is fused to a 2A self-cleaving peptide followed by, for example, GFP or blasticidin. This way, one can easily select a polyclonal population of high Cas9 expressing cells by using GFP or blasticidin, respectively, as a “selection” marker. This strategy is favored over the generation and subsequent validation of single cell derived clonal lines (18).

There is a general preconception in the CRISPR field that high Cas9 expression can result in toxicity and increasing off-target effects. Indeed, several reports in different model organisms have shown that high Cas9 expression is toxic (19). We also observed that mammalian cell lines expressing very high levels of Cas9 tend to downregulate Cas9 expression over time (data not shown). To address whether high Cas9 expression increases off-target effects, we analyzed the behavior of sgRNAs targeting safe-havens in our screens. We did not observe any increase in the number of outliers in the conditions with high Cas9 (Figs. 2B, 2C, 3C and SF 2D-E). Instead, the same outliers were found across the different levels of Cas9 expression and, as expected, became more pronounced in the high Cas9 conditions due to the faster editing speed. This indicates that off-targets are predominantly caused by poor sgRNA design and not by the level of Cas9 expression. However, because high Cas9 expression exacerbates sgRNA effects, off-target effects affecting cell fitness, do become more apparent.

## CONCLUSIONS

Our data highlight that Cas9 expression is a crucial parameter that influences the efficiency and timing of gene editing and its careful optimization per model system can significantly impact the outcome of high-throughput CRISPR screens. When establishing a CRISPR screening model, it is advisable to first generate a population of cells expressing sufficiently high levels of Cas9. Therefore, using a 2-vector system allowing for the selection of high Cas9-expressing population of cells and exclusion of low Cas9-expressing cells is preferred.

## METHODS

### Cell culture

MCF10A, SW480, A375 and HEK293T cell lines were obtained from ATCC. MCF10A cells were cultured in DMEM/F-12 medium containing 2.5 mM L-glutamine and 15 mM HEPES, supplemented with 5% horse serum, 10 µg/mL insulin, 0.5 µg/mL hydrocortisone and 0.1 µg/mL cholera toxin. SW480 cells were cultured in RPMI medium; A375 and HEK293T cells were cultured in DMEM medium. All the media were supplemented with 10% FBS, 1% penicillin/streptomycin and 2LmM L- glutamine. All cell lines were cultured at 37°C and with 5% CO_2_. All cell lines were validated by STR profiling and mycoplasma tests were performed every 2-3 months.

### Western blots

After the indicated culture period, cells were washed with chilled PBS, then lysed with RIPA buffer (25mM Tris-HCl, pH 7.6, 150 mM NaCl, 1% NP-40, 1% sodium deoxycholate, 0.1% SDS) containing Complete Protease Inhibitor cocktail (Roche) and phosphatase inhibitor cocktails II and III (Sigma). Samples were then centrifuged for 10 min at 15,000 x g at 4°C and supernatant was collected. Protein concentration of the samples was normalized after performing a Bicinchoninic Acid (BCA) assay (Pierce BCA, Thermo Scientific), according to the manufacturer’s instructions. Protein samples (denatured with DTT followed by 5 min heating at 95°C) were then loaded in a 4-12% polyacrylamide gel. Gels were run (SDS-PAGE) for approximately 45 min at 175 volts. Proteins were transferred from the gel to a polyvinylidene fluoride (PVDF) membrane at 330 mA for 90 min. After the transfer, membranes were incubated in blocking solution (5% bovine serum albumin (BSA) in PBS with 0.1% Tween-20 (PBS-T)). Subsequently, membranes were probed with primary antibody in blocking solution (1:1000) overnight at 4°C. Membranes were then washed 3 times for 10 min with PBS-T, followed by 1 h incubation at room temperature with the secondary antibody (HRP conjugated, 1:10,000) in blocking solution. Membranes were again washed 3 times for 10 min in PBS-T. Finally, a chemiluminescence substrate (ECL, Bio-Rad) was added to the membranes and signal imaged using the ChemiDoc-Touch (Bio-Rad).

### Generation of Cas9-expressing cancer cell lines

SW480 cells were transduced with Lenti-iCas9-neo (Addgene #85400) at approximately 60% confluence in the presence of polybrene (8 μg/mL). Cells were incubated overnight, followed by replacement of the lentivirus-containing medium with fresh medium containing G418 (100 μg/mL). After selection was completed, a titration of doxycycline was performed and the induction of Cas9 expression was assessed by flow cytometry. We tested a concentration range from 1 to 2000 ng/mL and observed a plateau of maximum expression above 200ng/mL and no toxicity up to 1000ng/mL. We choose 10, 40 and 1000 ng/mL of doxycycline for the induction of low, medium and high levels of Cas9 expression, respectively. Cas9 expression levels were confirmed by Western blot and flow cytometry one week later. The Cas9- expressing cell line was named “SW480_iCas9”.

SW480, A375 and HEK293T cells were transduced with lentiCas9-EGFP (Addgene #63592) at approximately 40-60% confluence in the presence of polybrene (8 μg/mL). Cells were incubated overnight, followed by replacement of the lentivirus-containing medium with fresh medium. After 1 week in culture cells were sorted on low, medium and high GFP levels (BD FACSAria™ Fusion Cell Sorter). One week later, Cas9 expression levels were confirmed by Western blot and flow cytometry. The Cas9-expressing cells were named according to their cell line name and Cas9 expression level, i.e. “*cell line name*_Cas9*^expression^ ^level^*”.

SW480 cells were also transduced with lentiCRISPR v2 (Addgene #52961) and with Lenti-Cas9-2A-Blast (Addgene #73310) at approximately 60% confluence in the presence of polybrene (8 μg/mL), using a MOI of 0,3 and >1, respectively. Cells were incubated overnight, followed by replacement of the lentivirus-containing medium with fresh medium containing puromycin (2 μg/mL) and Blasticidin (20 μg/mL), respectively. After the antibiotic selection was complete, Cas9 expression levels were confirmed by Western blot.

MCF10A cells were transduced with a lentivirus containing Edit-R Inducible Cas9 (Horizon CAS11229) at approximately 40% confluence in the presence of polybrene (4 μg/mL), using a MOI >1. Cells were incubated overnight, followed by replacement of the lentivirus-containing medium with fresh medium containing Blasticidin (10 μg/mL). After selection, several single cell clones were generated and Cas9 expression was assessed by Western blot. The clone with the highest Cas9 expression upon doxycycline treatment, and undetectable Cas9 expression in the absence of doxycycline (named “MCF10A_Edit-R”) was used for subsequent experiments.

MCF10A cells were also transduced with lentiCRISPR v2 at approximately 40% confluence in the presence of polybrene (4 μg/mL), using a MOI of 0,3. Cells were incubated overnight, followed by replacement of the lentivirus-containing medium with fresh medium containing puromycin (2 μg/mL). After the antibiotic selection was complete, Cas9 expression levels were confirmed by Western blot.

### Editing efficiency assessment by TIDE analysis

SW480_iCas9 cells were transduced with 3 different sgRNAs cloned into lentiGuide-Puro-T2A-GFP (see sgRNA cloning section below) at approximately 60% confluence in the presence of polybrene (8 μg/mL). Cells were incubated overnight, followed by replacement of the lentivirus-containing medium with fresh medium containing puromycin (2 μg/mL). Cells were kept in puromycin for 7 days. At day 7, a fraction of the cells was harvested, another fraction was analyzed by flow cytometry to confirm equal transduction efficiencies indicating similar sgRNA expression levels, and the rest of the cells were placed back in culture and treated with either 10 ng/mL or 1 µg/mL doxycycline. Cells were harvested from these two induction arms after 2, 4, 8, 12, 16, (20, 24 and 32 – only for sgRNA 3) days in continuous culture with doxycycline. DNA was isolated from all samples, Sanger sequencing was performed and editing efficiency was analyzed using TIDE (https://tide.nki.nl/). At day 13, cells were also harvested for western blot and flow cytometry analysis, to assess Cas9 expression levels.

### sgRNA cloning

sgRNAs targeting 3 different locations in the genome were cloned into a modified version of pU6-sgRNA EF1Alpha-puro-T2A-BFP (Addgene, #60955), where BFP was replaced by superfolder GFP (sfGFP) – named “lentiGuide-Puro-T2A-GFP”. Puro-T2A-BFP was removed using NheI and EcoRI sites. To introduce Puro-T2A- sfGFP, we amplified Puro-T2A as well as sfGFP, adding homology arms to both PCR products. Puromycin-T2A was amplified using the following oligos: FW: 5’- GTTTTTTTCTTCCATTTCAGGTGTCGTGAGCTAGCCCACCATGACCGAGTACAAGCCCAC-3’, RV: 5’- AACTCCAGTGAAAAGTTCTTCTCCTTTGCTGGTGGCGACCGGTGGGCCAGGAT TCTCCTC-3’ sfGFP was amplified using the following oligos: FW: 5’- GAGGAGAATCCTGGCCCACCGGTCGCCACCAGCAAAGGAGAAGAACTTTTCACTGGAGTT-3’ RV: 5’- ATGTATGCTATACGAAGTTATTAGGTCCCTCGACGAATTCTTATTTGTAGAGCTC ATCCA-3’

The resulting PCR products were inserted into the open sgRNA vector backbone through Gibson Assembly. To introduce the custom designed sgRNA sequences into the pU6-sgRNA-EF1-Puro-T2A-GFP vector, the vector was digested using BstXI and BamHI. The sgRNAs were PCR-amplified using sgRNA-specific forward primers and a universal reverse primer: FW_1: 5’- TTGGAGAACCACCTTGTTGGAATATGTTTAAGCCTAGAGAGTTTAAGAGCTAAGCTGGAA, FW_2: 5’- TTGGAGAACCACCTTGTTGGTATAGGATAATAGCTGGAAGGTTTAAGAGCTAAGCTGGAA, FW_3: 5’- TTGGAGAACCACCTTGTTGGAGAGGTCTAATTCTAGGGCCGTTTAAGAGCTAAGCTGGAA, RV: 5’- GTAATACGGTTATCCACGCGGCCGCCTAATGGATCCTAGTACTCGAGA.

The resulting PCR products were isolated and used for Gibson Assembly.

### Generation of custom sgRNA library

For the design of the custom sgRNA library targeting essential genes and safe-havens we used the Broad GPP sgRNA design portal and the safe-havens as designed previously (20). The sgRNA sequences (Supplemental Table 1) were ordered as a pool of oligonucleotides (Agilent) with flanking sequences to enable PCR amplification and Gibson assembly into lentiGuide-Puro (pLG, Addgene #52963). The pooled oligo library was amplified using pLG_U6_foward 5’- GGCTTTATATATCTTGTGGAAAGGACGAAACACCG-3’ and pLG-TRACR_reverse 5’-GACTAGCCTTATTTTAACTTGCTATTTCTAGCTCTAAAAC-3’. The fragments were purified and cloned into pLG as described by Morgens (21). The representation of the custom sgRNA library was validated by next generation sequencing.

### sgRNA libraries and screens

Two different versions of the Brunello library were used – a “1 vector system” (backbone expresses both Cas9 and the sgRNA library – Addgene #73179) and a “2 vector system” (backbone expresses only the sgRNA library – Addgene #73178). In this study we also used our Essential/Safe-havens library described above.

The appropriate number of cells to achieve 250-fold representation of the library, multiplied by five to account for 20% transduction efficiency, were transduced at approximately 40-60% confluence in the presence of polybrene (4-8 μg/mL) with the appropriate volume of the lentiviral-packaged sgRNA library. Cells were incubated overnight, followed by replacement of the lentivirus-containing medium with fresh medium containing puromycin (2-4 μg/mL). The lentivirus volume to achieve a MOI of 0,2 as well as the puromycin concentration to achieve a complete selection in 3 days was previously determined for each cell line. Transductions were performed in triplicate. After puromycin selection, cells were split into the indicated arms (for each arm, the appropriate number of cells to keep a 250-fold representation of the library was plated at approximately 10-20% confluence) and a T_0_ (reference) time point was harvested. Cells were maintained as indicated. In case a passage was required, cells were reseeded at the appropriate number to keep at least a 500-fold representation of the library. Cells (enough to keep at least a 500-fold representation of the library, to account for losses during DNA extraction) were collected when indicated, washed with PBS, pelleted and stored at -80°C until DNA extraction.

### DNA extraction, PCR amplification and Illumina sequencing

Genomic DNA (gDNA) was extracted (Zymo Research, D3024) from cell pellets according to the manufacturer’s instructions. For every sample, gDNA was quantified and the necessary amount of gDNA to maintain a 250-fold representation of the library was used for subsequent procedures (for this we assumed that each cell contains 6.6 pg genomic DNA). Each sample was divided over 50 μl PCR reactions (using a maximum of 1 µg gDNA per reaction) using barcoded forward primers to be able to deconvolute multiplexed samples after next generation sequencing. PCR mixture per reaction: 10 μl 5x HF Buffer, 1 μl 10 μM forward primer, 1 μl 10 μM reverse primer, 0.5 μl Phusion polymerase (Thermo Fisher, F-530XL), 1 μl 10mM dNTPs, adding H_2_O and template to 50 μl. Cycling conditions: 30 sec at 98°C, 20× (30 sec at 98°C, 30 sec at 60°C, 1 min at 72°C), 5 min at 72 °C. The products of all reactions from the same sample were pooled and 2 μl of this pool was used in a subsequent PCR reaction using primers containing adapters for next generation sequencing. The same cycling protocol was used, this time for 15 cycles. Next, PCR products were purified using the ISOLATE II PCR and Gel Kit (Bioline, BIO-52060) according to the manufacturer’s instructions. DNA concentrations were measured and, based on this, samples were equimolarly pooled and subjected to Illumina next generation sequencing (HiSeq 2500 High Output Mode, Single-Read, 65 bp). Mapped read-counts were subsequently used as input for the further analyses.

### Bioinformatics Analysis

For each CRISPR screen the sgRNA count data for each sample was normalized for sequence depth using the method described by DESeq2^3^ with the difference that the total value instead of the median of a sample was used. Because of the composition of the sgRNA library with a large fraction of sgRNAs targeting essential genes, the T1 samples were corrected by dividing with the median of T1/T0 ratios for the population of non-essential sgRNAs. For the genome-wide CRISPR screen comparing the efficiency of the 1-step and 2-step systems, a differential analysis was performed using DESeq2 (22). The output was sorted on the DESeq2 test statistic with the most depleted sgRNA at the top. We then used MAGeCK Robust Rank Algorithm to determine enrichment of sgRNAs targeting each gene (23). For the ROC curves, the output of these two analyses were filtered for 50 positive and 50 negative controls genes as described by Evers and colleagues (16). The Comparisons of the distribution of different groups of sgRNAs were performed using the Wilcoxon test.

### Reagents

Primary antibodies: Tubulin (Sigma, T9026) and Cas9 (Cell Signaling, 14697). Secondary antibody: Goat Anti-Mouse IgG (H + L)-HRP Conjugate (BioRad, 1706516).

## DECLARATIONS

### Availability of data and materials

All data and procedures used in this article are detailed in the Methods section. Any additional clarification or information regarding codes can be provided upon request.

### Authors’ Contributions

R.L.B., R.B. and R.M. supervised all research. J.N. and R.L.B. wrote the manuscript. J.N., K.J., C.L., L.K., M.D., B.M., D.V. and H.H. designed, performed and analyzed the experiments. All authors commented on the manuscript.

## Acknowledgements

We thank the NKI’s Flow Cytometry and Sequencing facilities.

## SUPPLEMENTARY DATA

**Supplementary data to Figure 1 (Suppl. Fig. 1):**
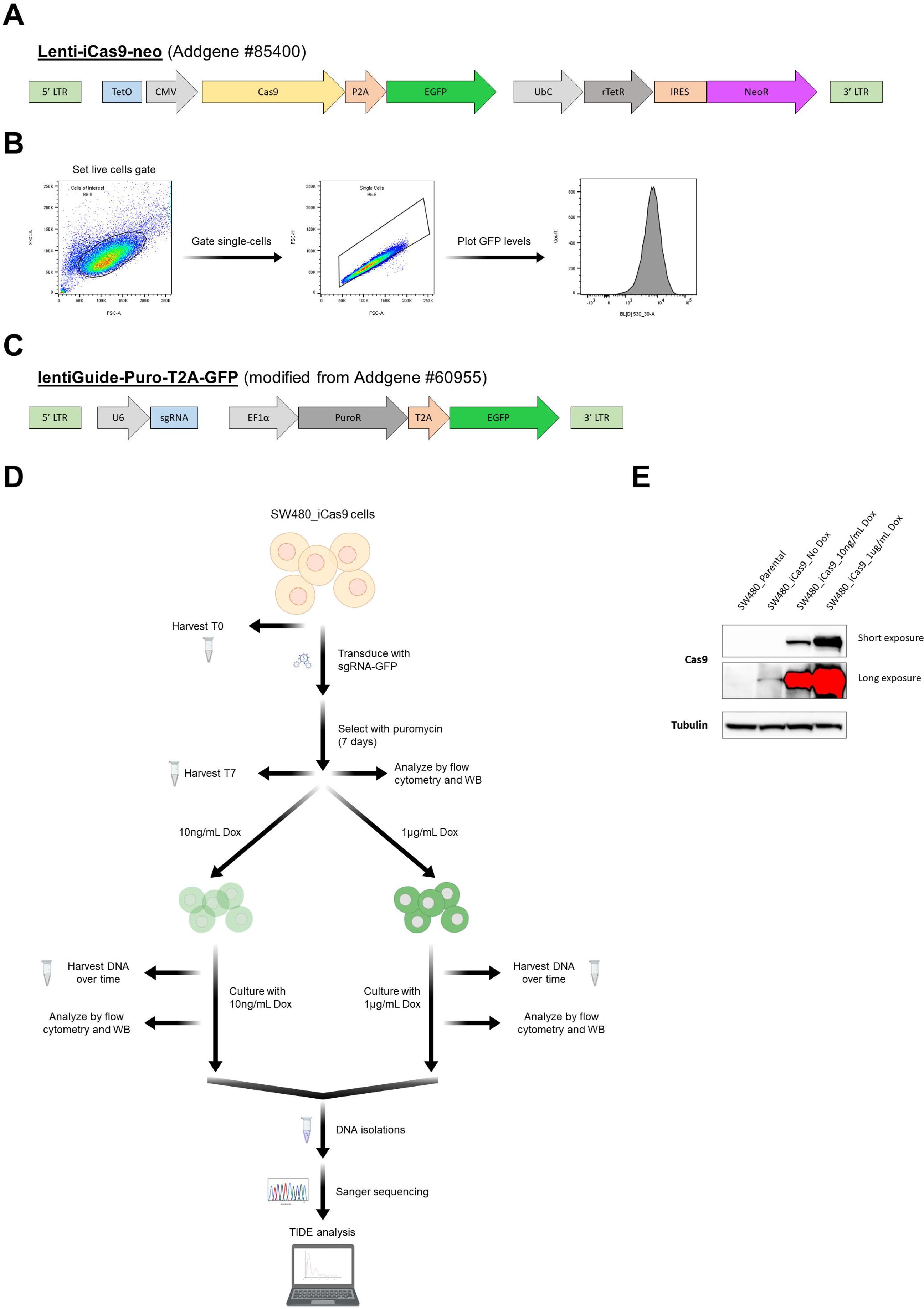
**A,** Schematic representation of the Lenti-iCas9-neo vector. **B,** Schematic representation of the gating strategy used for all flow cytometry experiments. Live cells were gated from all events, then single cells were gated from the live cells and finally the EGFP levels were plotted from the single cells in a histogram (y-axis = cell count, x-axis = EGFP levels). **C,** Schematic representation of the lentiGuide-Puro-T2A-GFP vector. **D,** Schematic of the experiment performed to assess editing efficiency by TIDE analysis. **E,** Analysis of the levels of Cas9 expression in parental SW480 cells and in SW480_iCas9 cells (using different concentrations of doxycycline) by western blot. Cells were harvested at day 13 of the experiment described in Fig. 1D. Tubulin was used as loading control.

**Supplementary data to Figure 2 (Suppl. Fig. 2):**
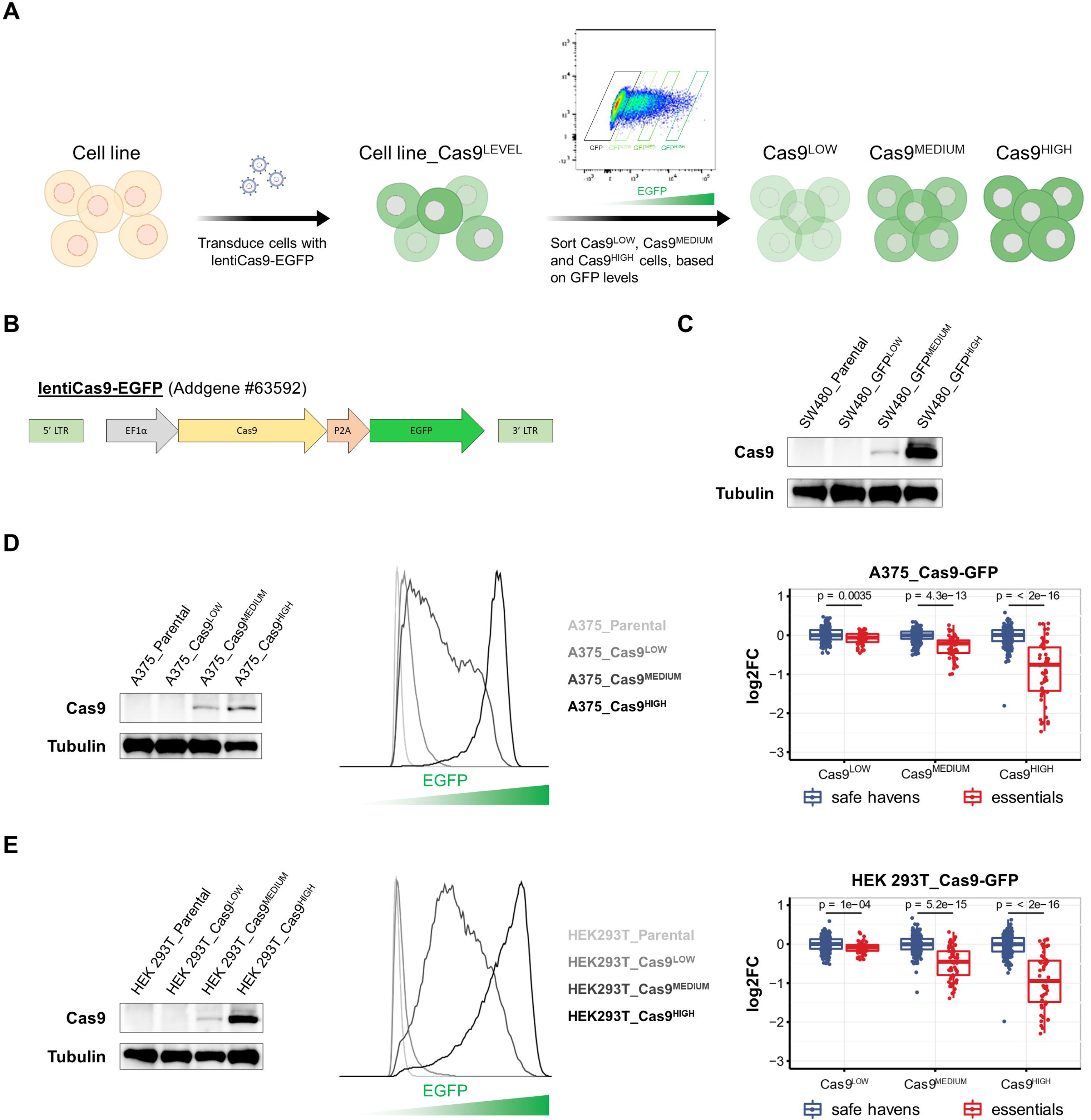
**A,** Schematic of the generation of cells lines with different Cas9 expression levels using the lentiCas9- EGFP vector. **B,** Schematic representation of the lentiCas9-EGFP vector. **C,** Analysis of the levels of Cas9 expression in SW480_Cas9-GFP cells by western blot. Tubulin was used as loading control. **D-E,** Performance of high-throughput screens is improved in conditions with higher Cas9 expression. In (D) analysis of the levels of Cas9 expression in A375_Cas9-GFP cells by western blot (left) and by flow cytometry (middle). On the (right) analysis of the of the depletion (log2 Fold Change) of the sgRNAs targeting essential genes and safe-havens in A375_Cas9-GFP cells with different Cas9 expression levels. Cells were screened with a library of sgRNAs targeting essential genes and safe-havens and cultured for approximately 8 population doublings. Box plot shows the median, interquartile range and the smallest and largest values no more than 1.5 times the interquartile range. Comparisons were made using the Wilcoxon test. In (E) analysis of the levels of Cas9 expression in HEK 293T_Cas9-GFP cells by western blot (left) and by flow cytometry (middle). On the (right) analysis of the of the depletion (log2 Fold Change) of the sgRNAs targeting essential genes and safe-havens in HEK 293T_Cas9-GFP cells with different Cas9 expression levels. Cells were screened with a library of sgRNAs targeting essential genes and safe-havens and cultured for approximately 8 population doublings. Box plot shows the median, interquartile range and the smallest and largest values no more than 1.5 times the interquartile range. Comparisons were made using the Wilcoxon test.

**Supplementary data to Figure 3 (Suppl. Fig. 3):**
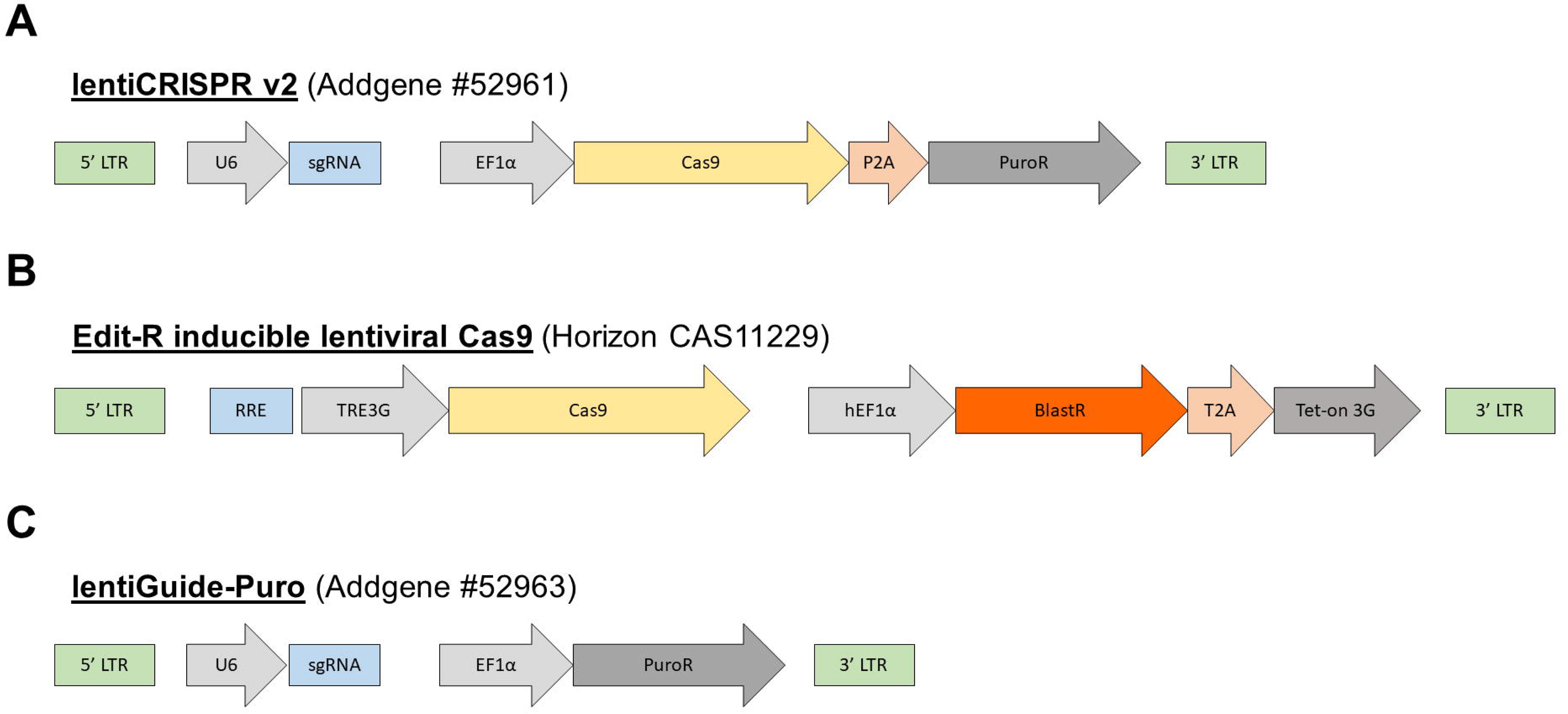
**A,** Schematic representation of the lentiCRISPR v2 vector. **B,** Schematic representation of the Edit-R inducible lentiviral Cas9 vector. **C,** Schematic representation of the lentiGuide-Puro vector.

**Supplementary Table 1:** Sequences of the sgRNAs present in the custom-made essential/safe-haven sgRNA library and results of the screens using this library.

**Supplementary Table 2:** Results of the depletion of the sgRNAs targeting essential and non-essential genes in MCF10A cells screened with the same sgRNA library (Brunello), using either a 1- step or a 2-step system.

